# Changing the Waddington landscape to control mesendoderm competence

**DOI:** 10.1101/867010

**Authors:** James R. Valcourt, Roya Huang, Sharmistha Kundu, Divya Venkatasubramanian, Robert E. Kingston, Sharad Ramanathan

## Abstract

As pluripotent human embryonic stem cells progress towards one germ layer fate, they lose the ability to adopt alternative fates. It is unknown how the cells’ competence for these alternative fates changes along their developmental trajectory or if this competence can be modulated. Here, we show that a differentiating stem cell’s probability of adopting a mesendodermal fate when given the appropriate signal falls sharply at a specific point along the ectodermal trajectory, and we further demonstrate that this point can be moved using genetic perturbations. Using a low-dimensional reaction coordinate to monitor progression towards ectoderm, we can determine the probability that individual cells at different points along this path can transition to the mesendodermal fate upon BMP4 and Activin A signal exposure. Knowing this probability allows us to prospectively isolate and profile differentiating cells based on their mesendoderm competence. Analysis and validation of these RNA-seq and ATAC-seq profiles identified transcription factors that can independently control the cell’s mesendoderm competence and its progression along the ectodermal developmental trajectory. In the classical picture of a Waddington landscape, these effects correspond to altering the barrier between fates and changing the cell’s location on the landscape, respectively. The ability of the underlying gene regulatory network to modulate these two aspects of the developmental landscape could allow separate control of the dynamics of differentiation and tissue size proportions.

## Main

Pluripotent cells have the ability to produce any of the myriad cell types seen in the adult body^1^, but they lose this potential as they differentiate. During initial lineage specification, cells can change their fate choice upon exposure to signals that induce an alternative selection^1–4^, such as by transplantation to a different location in the embryo. In time, however, the cell’s fate becomes determined, and it is no longer competent to choose a different lineage in response to the same external signals^5–7^. While lineage specification is relatively well-understood^8–17^, whether and how competence for adopting alternative lineages can be tuned during differentiation is not. Determining how this competence is set and modulated is essential for understanding developmental patterning and plasticity.

There is a fundamental challenge in understanding the competence of a cell to choose a specific fate in response to a signal because a cell’s fate choice is evident only after the expression of fate-specific marker genes. In mouse and humans, the fate markers for the germ layers are not expressed until at least twelve hours after exposure to the appropriate signals^20–22^. Since the gene expression and epigenetic state of the cell changes substantially during this period, determining how the molecular state of the cell at the time of signal exposure governs its competence to adopt alternative fates has been difficult.

To overcome this challenge, we use a framework motivated by the Waddington landscape^23^. In this picture, the pluripotent cell moves down into the valley corresponding to its chosen fate, and it is prevented from adopting the alternative fate by a barrier that rises between the valleys (Figure 1a). Thus, a cell’s ability to transition to the alternative fate depends both on its location along the developmental trajectory and the size of the barrier at that point. Using the germ layer selection of human embryonic stem cells (hESCs) as a model system, we measured the probability that a given cell can transition to the mesendoderm fate as a function of its position along the ectodermal developmental trajectory, thereby probing the location of the barrier between fates. Using this probability distribution, we could prospectively isolate and characterize cells based on their mesendoderm competence. Computational analysis of both the gene expression and epigenetic profiles of these populations allowed us to identify candidate genes that we predicted to control mesendoderm competence. By perturbing the levels of these genes, we were able to independently tune the barrier between fates and the cell’s position on the Waddington landscape. These dual mechanisms could play a role in regulating the proportions of the different germ layers in the developing embryo.

**Figure 1:**
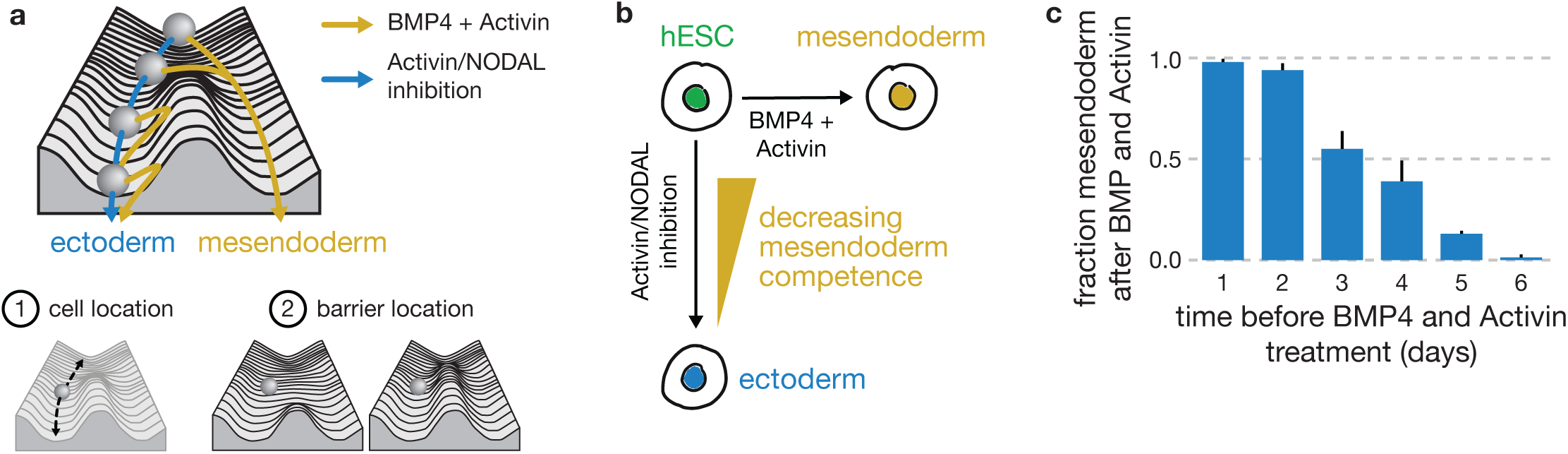
Stem cells lose competence to adopt mesendodermal fates upon BMP4 and Activin A signal exposure with increasing duration of Activin/NODAL inhibition. **a,** Schematic of a Waddington landscape illustrating the ectoderm (blue) / mesendoderm (yellow) fate choice. In this picture, the competence of a cell to produce mesendoderm depends both on the cell’s location along the ectodermal developmental trajectory and on the position of the barrier between the ectoderm and mesendoderm fates. **b,** hESCs choose the ectodermal lineage in response to Activin/NODAL inhibition and mesendodermal lineages in response to BMP4 + Activin A. **c,** Bar graph of the fraction of cells adopting mesendodermal fate as a function of the duration of ectodermal differentiation after which BMP4/Activin A was added (see Methods). At the population level, with an increasing duration of prior ectodermal differentiation, the competence to choose mesendodermal fates in response to signal decreases. Error bars: std (n=3).

## Results

### hESCs lose mesendoderm competence along the ectoderm developmental trajectory

Given the appropriate signals, hESCs *in vitro* can adopt either mesendodermal or ectodermal progenitor fates: BMP4 and Activin/NODAL signals induce mesendodermal fate^24^, while Activin/NODAL inhibition promotes ectodermal fate^21^ (Figure 1b). When cultured in conditions that allow the signaling ligands to access their corresponding receptors on the cells’ basolateral side (see Methods), hESCs exposed to BMP4 and Activin A uniformly produce cells expressing BRACHYURY, SOX17, and other markers associated with mesendoderm-derived cell types (Figure 1b, S1a, S1b). In contrast, inhibiting Activin/NODAL signaling promoted ectoderm-derived fates^21^, ultimately producing PAX6^+^ neurectoderm after 4-5 days (Figure 1b, S1c, S1d).

Consistent with previous data from mouse^22, 25^, we demonstrated that competence to adopt mesendoderm-derived fates decreases at the population level as cells differentiate towards the ectodermal fate. Increasing the duration of differentiation towards ectoderm reduced the fraction of cells that adopted mesendodermal fates in response to BMP4 and Activin A signal (Figure 1c, S1b, S1e, S1f). The temporal decrease in the mesendoderm fraction occurred despite the cells’ continued ability to respond to BMP and Activin signals throughout this period (Figure S1g, S1h). These findings suggested that the probability of cells transitioning to the mesendodermal fate in response to signal decreases over time within the differentiating population.

### Dynamics of OCT4 and SOX2 predict the mesendoderm competence of individual cells

We next sought to directly measure and predict the probability of individual cells adopting a mesendodermal fate in response to BMP and Activin signals along the ectodermal differentiation trajectory. To do so, we first developed a measure of each live cell’s position along the developmental trajectory by choosing a low-dimensional coordinate—in this case, the expression levels of key genes—whose dynamics accurately report on the choice of the two germ layer lineages. Our recent computational work allows us to identify these key genes for a given lineage decision from single cell gene expression data^26, 27^, and we have demonstrated the accuracy of this method for germ layer, neural, and hematopoietic development. For the decision between the two germ layer lineages, the genes that allow us to continuously monitor the progression of fate choice are the transcription factors OCT4 (POU5F1) and SOX2. Our previous work in mouse showed that the protein levels of Oct4 and Sox2 reflect the transitions of pluripotent cells to the mesendoderm or ectoderm fates^26–28^. We validated that, in humans as in mouse, both OCT4 and SOX2 are symmetrically highly expressed in the pluripotent stem cell, but they are asymmetrically downregulated in the two lineages. OCT4 expression is maintained in the mesendoderm while SOX2 is downregulated; in contrast, ectoderm differentiation involves SOX2 maintenance and OCT4 downregulation. Both TFs are also functionally important for these state transitions: OCT4 downregulation is necessary for neurectoderm induction^28, 29^, while SOX2 downregulation is required for mesendoderm fate selection^28^. Furthermore, direct conversion to a neural fate silences OCT4^28^, underscoring the fundamental nature of this reaction coordinate to the fate decision in question.

To monitor developmental trajectories in real time, we employed our validated hESC line in which one allele each of OCT4 and SOX2 had been replaced with OCT4:RFP and SOX2:YFP, respectively, at the endogenous locus^30^ (Figure S2a-c). Using flow cytometry, we followed the developmental trajectories of hESCs to ectodermal progenitors as they downregulated OCT4 over the course of six days. When BMP4 and Activin A signals were added at an intermediate stage of differentiation (Figure 2a), we could visualize a bifurcation of developmental trajectories: the mesendoderm-competent cells adopted OCT4^+^ SOX2^-^ mesendoderm-derived fates, while the cells that were not mesendoderm-competent proceeded towards OCT4^-^ SOX2^+^ ectoderm-derived fates.

**Figure 2:**
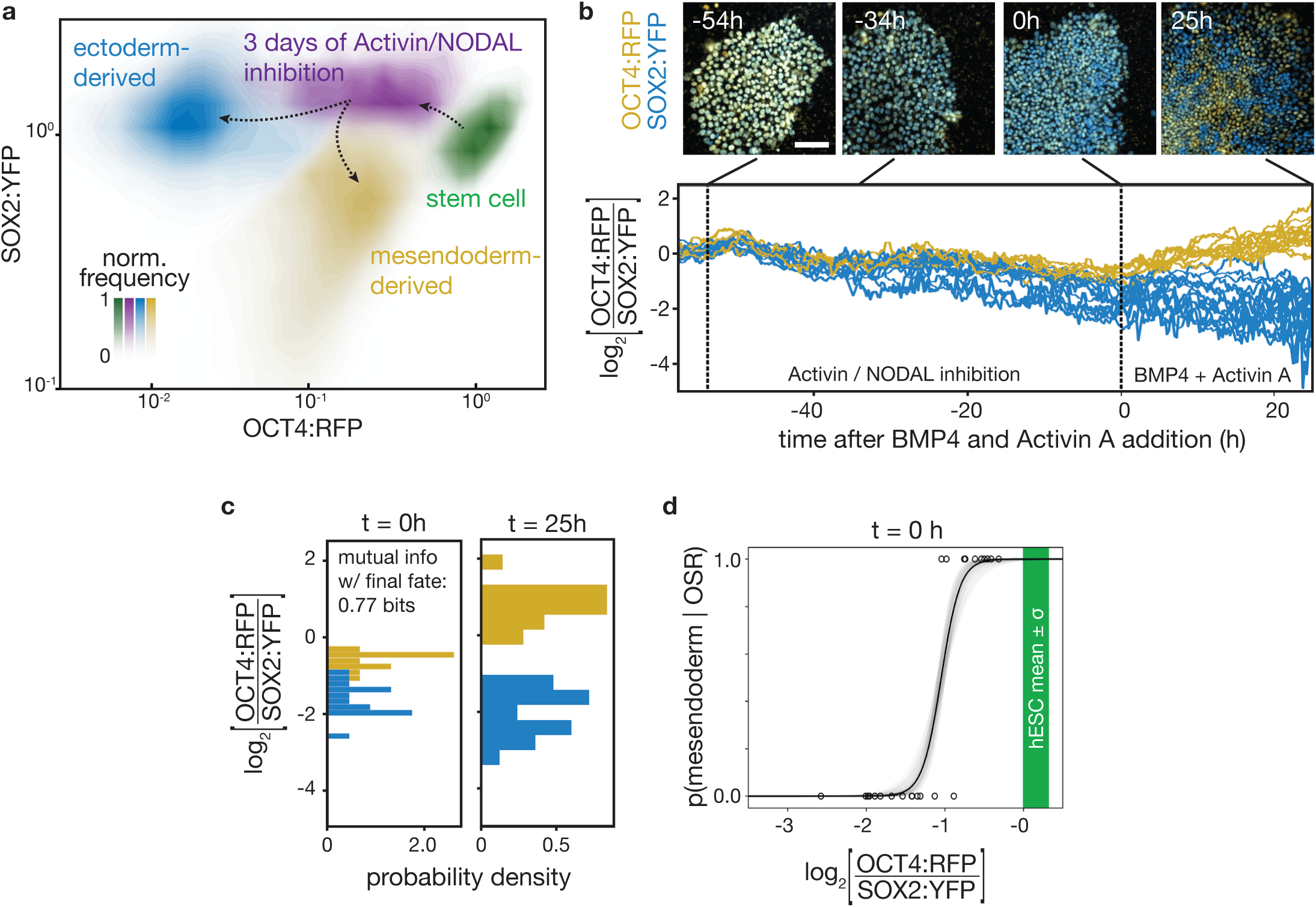
Dynamics of OCT4 and SOX2 accurately predict mesendoderm competence. **a,** Contour plots of flow cytometry data showing levels of OCT4:RFP and SOX2:YFP normalized to the mean level in the hESC population. hESCs (green) downregulate OCT4:RFP after 3 days of ectodermal differentiation (purple). After a subsequent two days of BMP4 + Activin A stimulation, the cells in this purple population bifurcate: the competent fraction chooses a mesendodermal fate (yellow), while the remaining continue to become ectoderm (blue). **b,** Top, snapshots from a time-lapse microscopy experiment of a field of hESCs showing endogenous OCT4:RFP (yellow) and SOX2:YFP (blue). Cells started in pluripotency conditions; at t=-54h, ectodermal differentiation began; at t=0h, BMP4 + Activin A stimulation began; at t=25h, experiment ended. Scale bar = 100 μm. Bottom, plot of the log ratio of the OCT4:RFP to SOX2:YFP signal (OSR) in individual cells through the time course above. Time traces of cells adopting a mesendodermal (ectodermal) fate at the end of the time course are shown in yellow (blue). **c,** Histogram of the log OSR from (b) shown at the time of signal addition (t=0h) and at the end of the experiment (t=25h). Histograms corresponding to cells adopting an eventual mesendodermal (ectodermal) fate at the end of the time course are in yellow (blue). The mutual information between the OSR at the moment of signal addition and the final fate choice is 0.77 bits. **d,** Plot of the probability of a single cell adopting a mesendoderm-derived fate given OSR, p(mesendoderm | OSR), calculated from time course in (b), mean black, 1 std in grey (see Methods). Green bar: mean value +/− std of the ratio in pluripotent stem cells.

We next sought to measure the probability of an individual cell adopting a mesendoderm-derived fate given the cell’s location along the ectodermal trajectory. To do so, we performed a time lapse experiment using the OCT4:RFP SOX2:YFP hESC line (Figure 2b). To monitor the cells throughout this process, we developed a custom live-cell microscopy setup that was capable of imaging cells on the flexible membrane every 15 minutes for over five days (Figure S2d). Based on the timing of mesendoderm competence loss in our flow cytometry experiments, we first differentiated the pluripotent stem cell population in this apparatus for 2.25 days in ectodermal differentiation conditions to obtain a heterogeneous population in which some cells had already lost mesendoderm competence and some had not. We then added BMP4 and Activin A signals for 25 hours, prompting mesendoderm-competent cells to adopt mesendoderm fates and non-mesendoderm-competent cells to adopt ectodermal fates.

Using our time lapse data, we demonstrated that each cell’s location along its developmental trajectory as measured by OCT4:RFP and SOX2:YFP levels was predictive of the cell’s mesendoderm competence, and that this measure allowed such prediction days before the expression of classical master regulators. We tracked individual cells from pluripotency through ectoderm-directed differentiation and subsequent BMP4 + Activin A signal (Figure 2b). Pluripotent cells were tightly localized in OCT4:RFP/SOX2:YFP space but downregulated OCT4:RFP at different rates upon ectoderm differentiation. Importantly, we found that, rather than the levels of the individual proteins, the OCT4:RFP to SOX2:YFP fluorescence ratio at the moment of BMP4 and Activin A signal addition was predictive of the ultimate fate of the cells with high accuracy (Figure 2c). Each cell carried 0.77 bits of mutual information (the maximum possible value being 1) about its mesendoderm competence state in its ratio of OCT4:RFP to SOX2:YFP at the moment of signal addition. Cells with a high ratio of OCT4:RFP to SOX2:YFP were able to become mesendoderm in response to the BMP4 and Activin A signal, while cells with a low ratio of OCT4:RFP to SOX2:YFP were not.

We then computed the probability of a cell adopting a mesendodermal fate given its OCT4:RFP to SOX2:YFP ratio (OSR) after normalizing OCT4:RFP and SOX2:YFP intensities to the mean levels measured in hESCs in pluripotency conditions (Figure 2d). The probability of adopting a mesendodermal fate given OSR, *p*(mesendoderm|OSR), had a sharp transition from 1 to 0, indicating that there was a defined point along the developmental trajectory at which cells lose their ability to become mesendoderm even when exposed to the relevant signals. On the Waddington landscape, the OSR corresponds to a cell’s position along the ectoderm trajectory, and the point at which *p*(mesendoderm|OSR) transitions from 1 to 0 corresponds to the location of the barrier that arises between the two fates.

### Prospective isolation of cell populations based on mesendoderm competence

Having computed *p*(mesendoderm|OSR) for single cells in our time lapse, we sought to predict the mesendoderm competence of cells in a heterogeneous differentiating population. Since cells in a population move along the developmental trajectory at differing rates, at any given time, *t*, the cells have a distribution of OSRs, *p*(OSR|*t*). In the Waddington landscape, this corresponds to the distribution of cell locations along the ectoderm trajectory at that given time. The fraction of the overall cell population that adopts a mesendoderm fate after BMP4 and Activin stimulation should be determined by the fraction of the cells that have not yet reached the point along the ectodermal trajectory where the barrier to mesendoderm arises. Thus, the total fraction of cells that become mesendoderm can be calculated by multiplying the fraction of cells with a given OSR by the probability that cells with that ratio will become mesendoderm, then summing over all OSRs (Figure 3a):

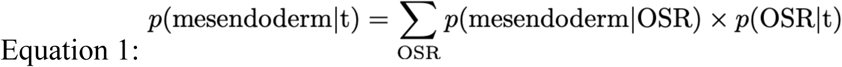

**Figure 3:**
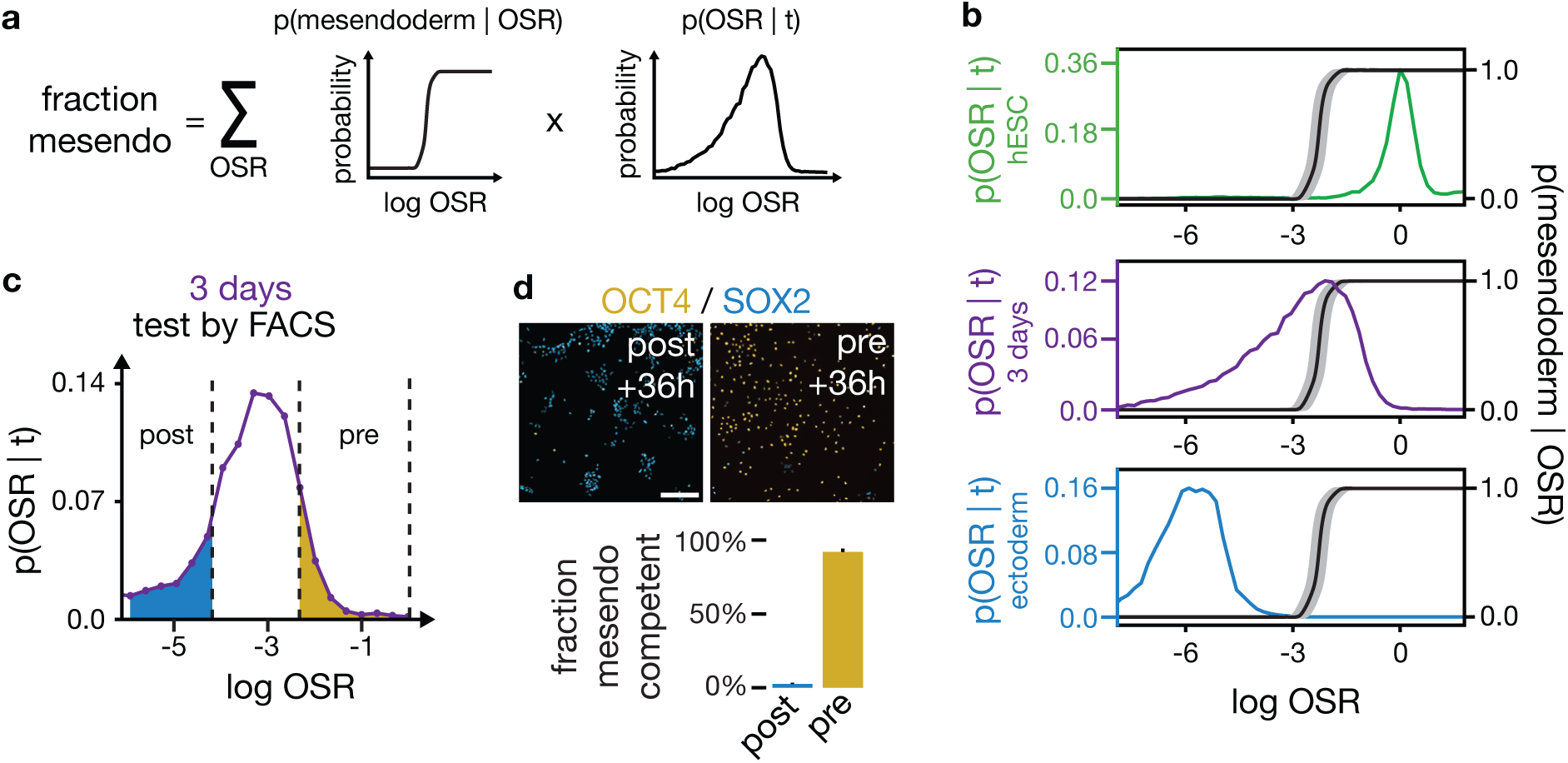
Prospective isolation of cell populations based on mesendoderm competence. **a,** The predicted fraction of cells that adopt a mesendodermal fate upon BMP4 and Activin stimulation at a given time t, p(mesendoderm | t), equals the sum over OSR of the probability of adopting a mesendodermal fate given OSR, p(mesendoderm | OSR), times the probability distribution of OSR in the population at that given time, p(OSR | t). **b,** Plot of p(OSR | t) as a function of OSR for the populations in (a) shown using the same color: pluripotent stem cells (green), 3 days of ectodermal differentiation (purple), ectodermal population (blue). p(mesendoderm | OSR) (black curve) inferred based on the final mesendoderm fraction from (a). p(OSR | t) moves leftward as cells differentiate. The cells in the purple population corresponding to the region with p(mesendoderm | OSR)=1 are predicted to be mesendoderm competent while the cells where p(mesendoderm | OSR)=0 are predicted not to be. **c,** Histogram showing p(OSR|t) for a cell population at t=3 days of Activin/NODAL inhibition. Overlaid on the histogram are the FACS gates based on (f) used to sort subpopulation “post,” shown in blue, predicted to have lost mesendoderm competence; and subpopulation “pre,” shown in yellow, predicted to retain mesendoderm competence. **d,** Top, images of pre- and post-competence-loss subpopulations after cell sorting by FACS followed by 40h of BMP4 + Activin A stimulation and immunostaining for OCT4 (yellow) and SOX2 (blue). Scale bar = 300 μm. Bottom, fraction of cells in the pre and post sorted population adopting mesendodermal fate. Error bars: std (n=3). The sorted populations essentially uniformly adopt the predicted fate.

We next reasoned that we should be able to use our ability to predict mesendoderm competence to prospectively isolate competent from non-competent cells from a single differentiating population. To this end, we measured *p*(OSR|*t*) of a population of differentiating hESCs using flow cytometry. Our analysis suggested that the cells in the high OSR region where our computed *p*(mesendoderm|OSR) ≈ 1 would be competent to differentiate into mesendoderm, while those in the low OSR region where *p*(mesendoderm|OSR) ≈ 0 would not (Figure 3b). We will hereafter refer to these populations as “pre-competence-loss” and “post-competence-loss,” respectively. To validate our predictions, we sorted cells based on their OSR using fluorescence-activated cell sorting (FACS) from a population that had been subjected to 3 days of ectodermal-directed differentiation (Figure 3c). We then added BMP4 and Activin A to the sorted populations to compare our predicted mesendodermal competence with the observed fate choice of these cells.

As predicted, we obtained essentially pure populations of OCT4^-^ SOX2^+^ ectoderm-derived fates from the post-competence-loss population and OCT4^+^ SOX2^-^ mesendoderm-derived fates from the pre-competence-loss population (Figure 3d). Thus, we were indeed able to prospectively isolate individual cells at points before and after loss of mesendodermal competence.

Having achieved our first goal of predicting mesendoderm competence, we turned to our second goal: to change this competence by modulating the Waddington landscape. We hypothesized that we could change the fraction of mesendoderm competent cells in one of two ways: (1) by altering the position of cells along the ectoderm-directed developmental trajectory, *p*(OSR|*t*), so that more or fewer cells would be past the point of competence loss at the time of BMP4 + Activin A signal addition, or (2) by modulating mesendoderm competence along the trajectory, *p*(mesendoderm|OSR), such that competence loss (the sharp drop-off in probability from 1 to 0) happens earlier or later along the trajectory. We further hypothesized that important factors controlling these two aspects would be DNA-binding factors whose expression patterns or access to binding sites changed along the developmental trajectory. Thus, to identify candidate factors that control these two probabilities, we characterized pre- and post-competence-loss populations using RNA sequencing (RNA-seq) and Assay for Transposase-Accessible Chromatin using sequencing (ATAC-seq).

### Key TF families are remodeled upon loss of mesendoderm competence

For our RNA-seq and ATAC-seq experiments, we isolated populations of pre- and post-competence cells using FACS from a single heterogeneous population of stem cells that had been subjected to 3 days of ectodermal differentiation. As a control, we reserved a small fraction of sorted cells from each sample that we then treated with BMP4 and Activin A to confirm the competence of that sorted population. We also included a third, mesendoderm-derived population produced by subjecting pluripotent stem cells to BMP4 + Activin A for 40 hours, which allowed us to distinguish lineage-specific changes in expression and chromatin accessibility from changes that are shared by cells entering either lineage (gFigure S3a). These populations displayed significant and concerted changes in both their RNA-seq and ATAC-seq signatures upon the loss of mesendoderm competence (Figure 4a).

**Figure 4:**
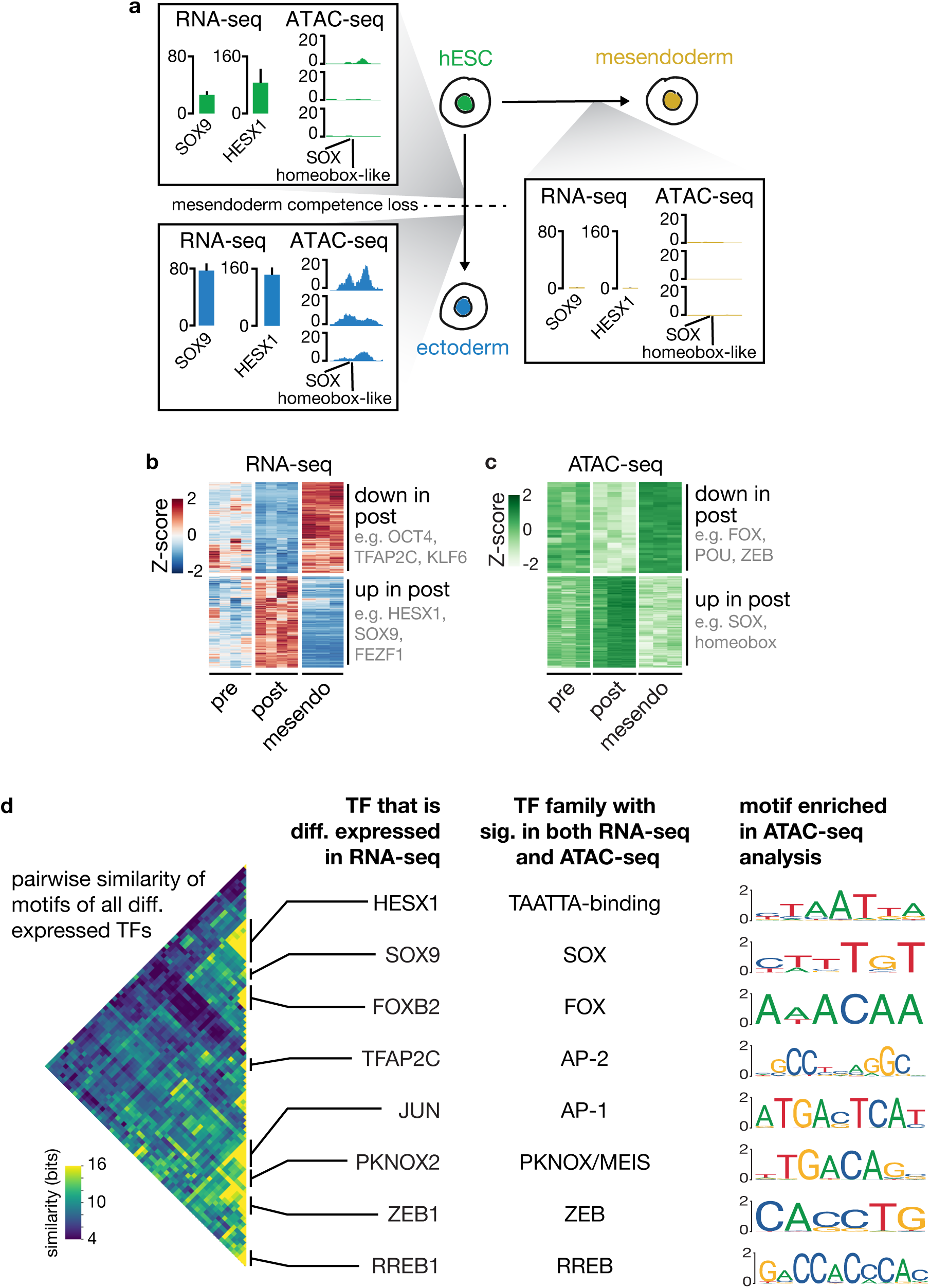
Key TF families show concordant changes in expression and motif accessibility upon mesendoderm competence loss, suggesting perturbation candidates. **a,** Illustration showing the three assayed populations: pre- and post-competence loss (Left) and mesendoderm (Right). Inset boxes illustrate, with one example, combined analysis of gene expression and chromatin accessibility data from the three populations. Within each inset: Left, gene expression of SOX9 and HESX1 measured in transcripts per million, error bars: sd (n=3). Right, ATAC-seq read depth for three biological replicates at a genomic locus that contains SOX and TAATTA homeobox-like motifs, to which SOX9 and HESX1, respectively, are known to bind. SOX9 and HESX1 are upregulated in the post-committed population along with increased chromatin access to their binding site. **b,** Shown are all genes with significant lineage-specific differential expression level changes (Heat map with Z-scores) between the pre-competence-loss and post-competence loss populations (n=4) as well as mesendoderm-derived outgroup (n=3 biological replicates). Key TFs downregulated (upregulated) upon loss of mesendoderm competence include OCT4, TFAP2C, and KLF6 (HESX1, SOX9, and FEZF1). **c,** Heatmap showing row-normalized ATAC-seq read depth in all 250 bp peaks with a significant change in read depth between competent and non-competent populations (n=3). Like gene expression, these regions display clear, commitment and lineage-specific accessibility changes. **d,** Left, heatmap showing an information-based measure of similarity (see Methods) between the known DNA binding motifs of all pairs of differentially expressed TFs. Each row/column corresponds to the motif of one TF, and the matrix is arranged using hierarchical clustering. Only one half of the symmetric matrix is shown. TFs with similar preferences for DNA primary sequence cluster together, and notable families are labeled with the name of one cluster member in the column at Center Left. Center Right, the name of the TF family whose signatures are seen in both the RNA-seq analysis and ATAC-seq analysis. Right, the corresponding motif identified as significant in the ATAC-seq analysis for each labeled TF family. These key TF families create concordant signatures in gene expression and chromatin accessibility data during mesendoderm competence loss, and are good candidates for perturbing mesendoderm competence.

Differential analysis of our RNA-seq data between the pre- and post-competence-loss populations using mesendoderm as an outgroup (see methods) showed that 544 genes were upregulated specifically in the post-competence-loss cells, 23 of which were TFs (Figure 4b, S3b). We also found 673 genes (32 TFs) that were specifically downregulated. In particular, we observed the differential expression of TFs such as SOX9, HESX1, LHX2, FOXB2, TFAP2A, TFAP2C, PKNOX2, ZEB1, ZEB2, and GBX2, along with the expected expression pattern of OCT4 (Figure S3c). Consistent with our earlier observation (Figure S1g-h), the differentially expressed genes did not include signaling pathway components (Figure S3d). Our data further showed that competence loss occurred prior to the expression of master regulators such as PAX6 and SOX1 (Figure S3c). The expression pattern of all TFs is plotted in Figure S3e.

We validated our findings with complementary analysis of GO term enrichment (Figure S3f) and ChIP-seq target enrichment (Figure S3g). Again, we observed that genes that are specifically downregulated upon competence loss are most enriched for putative targets of factors previously implicated in pluripotency and germ layer selection, such as SOX2, ZEB1, SMAD4, TCF3, and KLF4; in contrast, genes that are specifically upregulated upon mesendoderm competence loss are most enriched for putative targets of Polycomb repressive complex 2 component SUZ12, which is known to repress ectoderm target genes until their expression is appropriate.

From our ATAC-seq analysis, we observed DNA accessibility peaks that showed reproducible, clear changes between groups, alongside many peaks that were present in all samples (Figure S4a). Accessibility, as assayed by read depth, showed clear peaks at transcription start sites (Figure S4b). Differential analysis of our ATAC-seq data between the pre- and post-competence-loss populations using mesendoderm as an outgroup (see Methods) showed thousands of regions that change accessibility between pre- and post-competence-loss populations. We found 2071 regions that were more accessible after competence loss, and 233 that were less accessible (FDR < 0.05; Figure 4c). Most of these differentially accessible regions were located in distal intergenic or intronic regions (Figure S4c).

We next confirmed that we could reproduce expected patterns in our ATAC-seq data. Using the software GREAT^31^, we found that genes near loci with increased accessibility in the post-competence-loss samples were most significantly enriched for orthologs of mouse genes expressed in the Theiler Stage 11 neurectoderm (Figure S4d), increasing our confidence that we were indeed sampling the transition we hoped to capture. We further noted that we could recover previously hypothesized motif signals from our data. For example, our ATAC-seq peaks that changed significantly and which also contained a compressed OCT4/SOX17 motif were almost exclusively open in the mesendoderm-derived outgroup (Figure S4e), consistent with what would be predicted from previous work^32^; in contrast, such regions that contained the classical OCT4/SOX2 motif were generally closed in the mesendoderm-derived cells. Interestingly, we did not observe significant changes in accessibility at any of the ENCODE-annotated candidate regulatory elements of pluripotency genes such as OCT4, SOX2, NANOG, KLF4, and MYC (Figure S4f).

To find TFs that potentially bind to the differentially accessible regions, we searched for sequence motifs that were enriched at these loci. We found more than 20 such motifs, many of which matched the known DNA-binding motifs of the differentially expressed TF families we had identified, including motifs that were similar to those bound by SOX, forkhead box (FOX), AP-2, AP-1, TAATTA-binding homeobox-like, PKNOX/MEIS, Zinc Finger E-Box Binding Homeobox (ZEB), and POU family TFs (Figure S4g), the latter of which includes OCT4 as a member. As a complementary analysis, we determined which known sequence motifs could best explain the changes in chromatin accessibility that we observed across the point of competence loss (Figure S4h).

We clustered the known binding motifs of the differentially expressed TFs by calculating a measure of similarity between each pair of motifs, and we found that many such TFs shared similar binding motifs (Figure 4d). These clusters correspond to major TF families—including the FOX, SOX, AP-2, PKNOX/MEIS, ZEB, and homeobox-like TAATTA-binding TF families—and multiple members of each family are differentially expressed. Each of these major TF family DNA binding motifs was also enriched in the ATAC-seq analysis, indicating that the expression changes of these TFs have clear signatures in the chromatin accessibility data. Taken together, the gene expression and chromatin accessibility data revealed that a small, core set of TFs from key families are remodeled upon loss of mesendodermal competence (Figure S5a).

### Key TFs independently modulate barrier between fates and location along developmental trajectory

We hypothesized that perturbation of these factors could modulate the position of cells along the ectoderm-directed developmental trajectory, *p*(OSR|*t*), or the cell’s mesendoderm competence along the trajectory, *p*(mesendoderm|OSR). To test these predictions, we selected 36 genes, composed largely of the differentially expressed TFs in our core network plus their paralogs (Figure S5b and S5c; see Methods), for overexpression studies. We seeded two parallel samples of pluripotent stem cell populations for each candidate gene. Using a lentiviral delivery system, we transduced cells in both samples with a payload of a single gene of interest separated from the C-terminal end of a mCerulean cyan fluorescent protein (CFP) by a P2A self-cleaving peptide sequence, all under the control of an EF-1α promoter (Figure S6a). Viral transduction was initiated at the onset of ectoderm-directed differentiation (Figure S6b). After three days of differentiation, we analyzed one sample by flow cytometry to measure *p*(OSR|*t*). We switched the other sample to media containing BMP4 + Activin A for 42 hours and assayed the final fate fractions by flow cytometry. We confirmed that our non-transduced controls had produced about 50% OCT4:RFP^+^ SOX2:YFP^-^ mesendoderm-derived cells and 50% OCT4:RFP^-^ SOX2:YFP^+^ ectoderm-derived cells. The transduced cells, in contrast, expressed CFP (Figure S6c) and displayed varying final lineage proportions. The co-expressed CFP marker allowed us to monitor transduction efficiency at the single-cell level. We titrated viral concentration to achieve <50% transduction so that each sample also contained many non-transduced (CFP^-^) cells to serve as an internal control population.

In principle, a change in the fraction of lentiviral-perturbed cells that form mesendoderm-derived cell types after BMP4 and Activin A treatment could result from either of the hypothesized effects: first, the perturbation could facilitate or impede the initial ectoderm-directed differentiation, thereby changing *p*(OSR|*t*); second, the perturbation could change *p*(mesendoderm|OSR), the probability of transitioning to mesendoderm given the location along the developmental trajectory. For each candidate TF, we computed *p*(mesendoderm|OSR) based on the *p*(OSR|*t*) and the observed final fraction of mesendoderm of lentiviral-perturbed cells (Equation 1).

For certain candidate TFs, their overexpression principally affected mesendoderm competence by shifting the barrier between fates, captured by *p*(mesendoderm|OSR). For example, overexpression of JUNB, POU2F3, or FOXB2 increased mesendoderm competence by shifting *p*(mesendoderm|OSR) (Figure 5a). We concluded that these candidates tuned mesendoderm competence by moving the barrier between fates on the developmental landscape.

**Figure 5:**
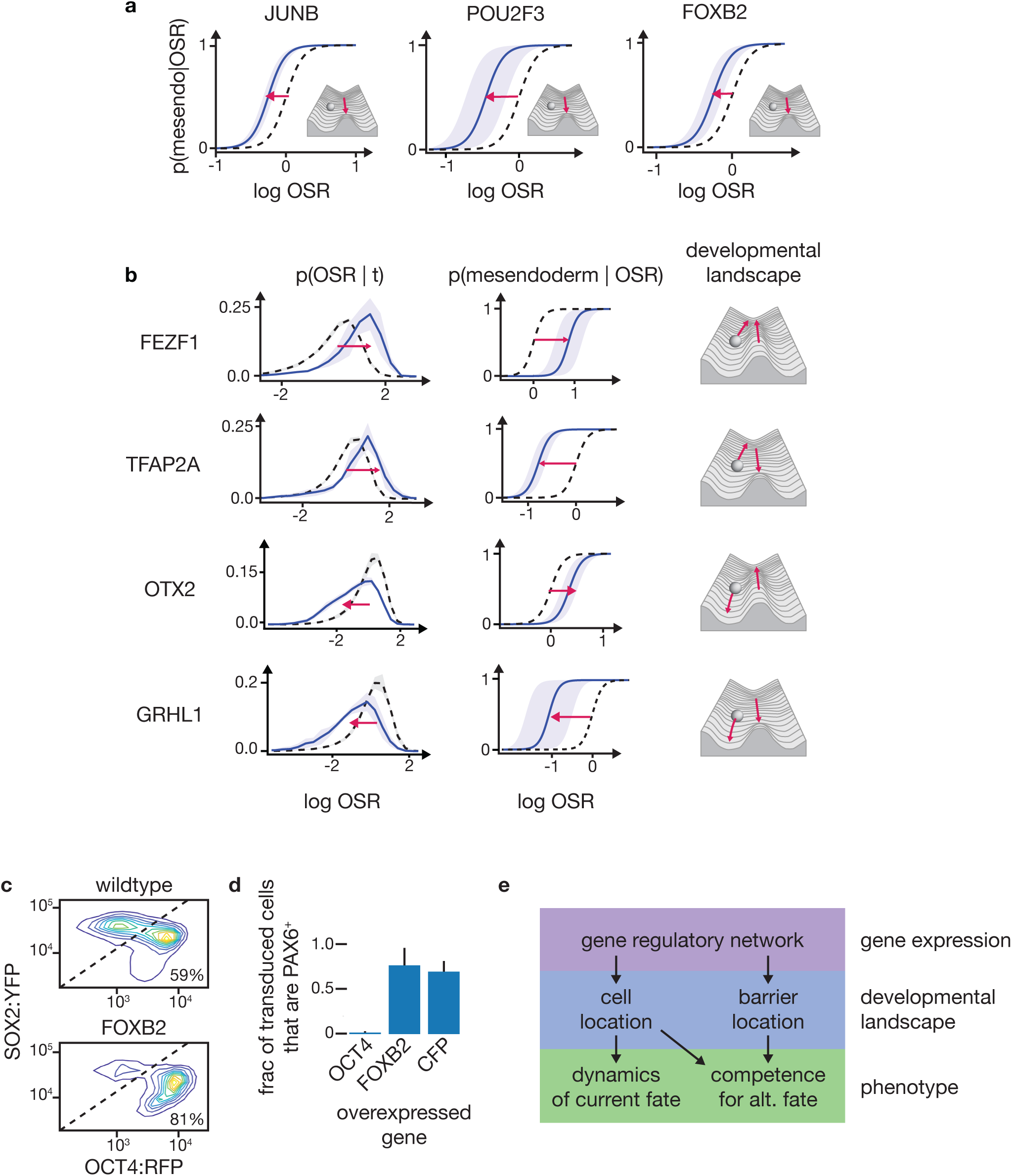
Perturbation of TFs candidates from RNA-seq and ATAC-seq analysis can independently tune progression along the developmental trajectory and inter-fate barrier. **a,** Over expression of three candidates, JUNB, POU2F3, and FOXB2, shifted p(mesendoderm | OSR) to keep cells mesendoderm competent for longer along the ectodermal differentiation trajectory. Black dotted: wildtype; blue: transduced cells; shaded: std (n=3, FOXB2 n=4). Inset, schematic depicting the effect of overexpression on the Waddington landscape. **b,** Over expression of FEZF1, TFAP2A, OTX2, and GRHL1 (top to bottom) altered both p(OSR | t) (left column) and p(mesendoderm | OSR) (middle column). Black dotted: wildtype control; blue: transduced cells, shaded: std (n=3). Left, p(OSR | t). Center, p(mesendoderm | OSR). Right, schematic as in (a). Overexpression of candidates can tune p(OSR | t) and p(mesendoderm | OSR) independently. **c,** Top, FACS density plot (OCT4 vs SOX2) of wild type cells (un-transduced CFP-) showing two peaks corresponding to ectodermal lineage above diagonal (41% of cells) and mesendodermal lineage, below diagonal (59% of cells). Bottom, transduced cells (CFP+) from the same population with CFP:P2A:FOXB2 cassette show 81% of cells adopting mesendodermal fate. Overexpression of FOXB2 increases the fraction of cells that adopt a mesendodermal fate. **d,** Fraction of cells adopting ectodermal fate (PAX6+) under ectodermal differentiation conditions. Cells with CFP:P2A:FOXB2 or the control CFP:P2A:CFP show normal ectodermal differentiation. OCT4 over expression precludes PAX6 expression and ectodermal differentiation. Error bars: std. **e,** The gene regulatory network governs both the progression of the cell along the developmental trajectory and shape of barrier between fates. In turn, both the cell’s location on the developmental landscape and the location of the barrier determine the cell’s fate in response to alternative signals. Only perturbations of the location of the barrier alters competence without altering the dynamics of movement along the original trajectory in the absence of a new signal.

In contrast, other candidates affected progression along the developmental trajectory, captured by *p*(OSR|*t*). For example, overexpression of SOX9 facilitated movement along the developmental trajectory, thereby shifting the distribution of *p*(OSR|*t*) towards the ectoderm fates (Figure S6d). In contrast, overexpression of TFAP2C hindered movement along the developmental trajectory, thereby shifting *p*(OSR|*t*) towards the pluripotent state. We concluded that these candidates tuned fate competence by changing cellular position along the developmental trajectory.

Notably, our candidates impacted competence by tuning *p*(mesendoderm|OSR) and *p*(OSR|*t*) independently of one another (Figure 5b, S6e). For example, while overexpression of FEZF1 and TFAP2A both shifted *p*(OSR|*t*) towards the pluripotent state, they shifted *p*(mesendoderm|OSR) in opposite directions. The outcomes seen upon overexpression of FEZF1, TFAP2A, OTX2 and GRHL1 together represent all four possible combinations of directional shifts in *p*(OSR|*t*) and *p*(mesendoderm|OSR). Thus, our candidates independently tuned *p*(OSR|*t*) and *p*(mesendoderm|OSR), thereby separately altering cellular location on the Waddington landscape and the barrier between fates.

Lastly, we explored the downstream developmental consequences of mesendoderm competence modulation via these two mechanisms. Overexpression of FOXB2, a candidate that increased mesendoderm competence primarily via a shift in *p*(mesendoderm|OSR), increased the fraction of cells that adopted mesendoderm by 32±5% (Figure 5c). Despite this change in mesendoderm competence, FOXB2 overexpression did not prevent normal neurectodermal differentiation as assayed by PAX6 expression in the absence of the BMP and Activin signal (Figure 5d, S6f). Thus, alternative fate competence can be modulated without preventing normal lineage progression in the absence of alternative-fate-inducing signals.

## Discussion

Our finding that competence for an alternative fate can be controlled either by changing the location of the cell along the developmental trajectory or moving the barrier that prevents cells from crossing over to the mesendoderm valley suggests possible evolutionary and developmental consequences. While moving the cell along the trajectory interferes with normal development in the absence of alternative signal, moving the barrier does not (Figure 5e). These two effects represent fundamentally different mechanisms for tuning competence, and both could be at play in the developing embryo. During the patterning of the mammalian epiblast, for example, the mesendodermal progenitors are generated along the primitive streak as it extends anteriorly from the posterior end of the epiblast. We speculate that changing the dynamics of epiblast competence loss anterior to the primitive streak could be a possible mechanism for tuning the length and extent of the streak. Further investigation of the role of competence modulation during mammalian gastrulation could be an important element in a full description of this important process, and the same mechanism could be acting to tune relative tissue sizes during any given cellular decision.

We expect that discovering reaction coordinates for different lineage decisions and monitoring the dynamics of differentiation along these coordinates will be fruitful for dissecting many cell fate decisions and state transitions. Our recent computational work has demonstrated how to find candidate genes that accurately reflect any given lineage choice^26, 27^, which greatly simplifies the prospect of following this approach in another context. Indeed, several other choices and competence restrictions occur immediately adjacent to the loss of mesendoderm competence in the early germ layer lineage tree, such as the presumptive loss of non-neural potential during neurectoderm fate determination or the loss of ectoderm competence during mesendoderm differentiation. In sum, these findings reveal two independently tunable mechanisms for adopting alternative fates along a developmental trajectory. Understanding how competence is tuned along distinct mechanistic axes will be crucial for elucidating the dynamics of mammalian embryonic patterning during development.

## Methods

### Cell lines

We conducted our experiments using WA01 (H1, WiCell) human embryonic stem cells. We also used an H1 cell line in which both OCT4 and SOX2 were tagged with fluorescent proteins as previously described^30^. In these cells, one endogenous copy of OCT4 was replaced with OCT4:tdTomato followed by an internal ribosomal entry site and a neomycin resistance gene to allow for selection, and one endogenous copy of SOX2 was replaced with SOX2:FLAG:Citrine:P2A:PuroR.

### Cell culture

hESCs were cultured in 6-well tissue culture dishes treated with Matrigel (Corning) and supplied with mTeSR media (STEMCELL Technologies) according to the manufacturer’s specifications. For routine culture, we passaged by washing with phosphate buffered saline (PBS) followed by ReLeSR (Stem Cell) treatment according to the manufacturer’s instructions. Cells were passaged in clumps of 8-10 cells and seeded in mTeSR supplemented with the Rho-associated protein kinase inhibitor γ-27632 (STEMCELL Technologies) at 10 µM for the first day to improve survival. All cell lines used were routinely tested for mycoplasma contamination.

For all experiments, we seeded cells on polyester membrane filters (Sterlitech) with 3 µm pores that had been treated with Matrigel. We chose this substrate to allow all cells to receive the BMP and Activin/NODAL signals we added to the media. TGF-β superfamily receptors, such as those for BMP4 and Activin A, are localized basolaterally in epithelial stem cell colonies and *in vivo* in the epiblast, so they are insulated from ligands in the apical media or luminal fluid^30, 33^. Typical tissue culture conditions allow for only the cells on the edge of the colony to receive signals, but growing cells on a membrane allows all cells in a colony access to the BMP and Activin ligands.

For live cell imaging, membranes were first glued to a custom 300 µm thick stainless-steel washer with Cytoseal 60 (Thermo Fisher), allowed to dry, sterilized with washes in 70% ethanol and with UV treatment, then treated with Matrigel for cell seeding.

### Differentiation conditions

Differentiation towards the ectoderm lineage was effected using mTeSR supplemented with 0.5 µM A83-01 (R&D Systems), a small molecule inhibitor of Activin and Nodal signaling. BMP4 + Activin A treatment was accomplished by treating cells with mTeSR supplemented with 3 ng/mL recombinant human BMP4 protein (R&D Systems) and 100 ng/mL recombinant human Activin A protein (R&D Systems). For neurectoderm-directed differentiation, we inhibited BMP signaling with 0.5 µM LDN-193189 in addition to Activin/Nodal inhibition with 0.5 µM A83-01 for 6 days.

### Flow cytometry

Cells were washed with PBS (Lonza) and removed from membranes by treatment with Accutase (Innovative Cell Technologies) until the cells were dissociated, about 20 minutes. Cells were analyzed on an LSRFortessa (BD Biosciences).

### Fluorescence activated cell sorting

Accutase-dissociated cells were sorted using a BD Aria III (BD Biosciences) using a 100 µm nozzle. Cells were gated such that the pre-competence-loss population was taken as the cells with the top 10-15% OCT4:RFP to SOX2:YFP ratio, and the post-competence-loss population was the bottom 10-15% OCT4:RFP to SOX2:YFP ratio. We sorted around 250,000 cells per subpopulation in a typical experiment. Populations were sorted into 1.5 mL centrifuge tubes (Eppendorf) filled with 500 µL of mTeSR supplemented with 10 µM γ-27632; by the end of the sort, ∼800 µL of sheath and sorted cells had been added to each tube. After the sort had completed, we pelleted the cells in a microcentrifuge at 250 xg for 3 minutes, then resuspended in PBS.

For each sorted sample, about 10% of the sorted cells were reserved for competence testing to confirm the pre-/post-competence-loss status of the sorted population. These cells were seeded back into glass-bottom 24-well plates (Ibidi) treated with Matrigel and filled with 1 mL of mTeSR supplemented with γ-27632 and allowed to recover for 3 hours. The media was then changed to mTeSR supplemented with BMP4 and Activin A for 36 hours. Cells were fixed and stained for OCT4 and SOX2 according to the protocols described under “Immunofluorescence.”

### RNA-seq

Total RNA was prepared from sorted or dissociated cells with an RNeasy Mini Kit (Qiagen) according to the manufacturer’s instructions. For the mesendoderm-derived outgroup samples, the input to the RNA extraction kit was a cell population directly after dissociation with Accutase; for FACS sorted populations, the input was sorted cells suspended in PBS. RNA integrity was quantified with a TapeStation 4200 (Agilent). All RINe scores were ≥ 9.9. Sequencing libraries were prepared by the Bauer Core at Harvard University using a Kapa mRNA Hyper Prep kit with Poly-A selection. Sequencing was performed on a NextSeq High output flow cell that generated paired-end 38bp reads. We obtained ≥42M reads per sample.

### ATAC-seq

ATAC-seq was performed as previously described^34^. Briefly, live cells were lysed and incubated with Tn5 transposase for 30 min at 37°C. After DNA purification, samples were amplified for the appropriate number of cycles as determined by qPCR to minimize PCR bias. Sequencing was performed by the sequencing core at Massachusetts General Hospital. We obtained ∼100M mapped paired-end reads per sample.

### Plasmid construction

Overexpression targets were subcloned from plasmids available through the Harvard PlasmID database, where available. Other targets were cloned from complementary DNA (cDNA) libraries. To prepare cDNA libraries for cloning, we differentiated human stem cells for 3 days in mTeSR + 0.5 µM A8301 and extracted RNA with RNeasy Mini Kit (Qiagen) according to the manufacturer’s instructions. We then performed first strand cDNA synthesis using SuperScript II Reverse Transcriptase (Thermo Fisher). We amplified the relevant cDNAs using Phusion polymerase (NEB) or Kapa HiFi (Kapa Biosystems). The OCT4 DNA binding domain and the SOX2 DNA binding domain (OCT4DBD and SOX2DBD) were amplified from cDNA. The OCT4DBD consisted of amino acids 131-296 of OCT4A (NCBI reference sequence: NM_002701.5). The SOX2DBD consisted of amino acids 37-117 of SOX2 (NCBI reference sequence: NM_003106.3). All cDNA-amplified clones were fully sequence confirmed by Sanger sequencing (Genewiz).

We constructed a vector from the second-generation lentiviral transfer backbone pWPXL with an EF-1α promoter. pWPXL was a gift from Didier Trono (Addgene plasmid #12257; http://n2t.net/addgene:12257; RRID:Addgene_12257). We first joined sequences for fluorescent protein mCerulean (CFP) and 2A peptide P2A (a ribosomal skip sequence) with Q5 (NEB) fusion PCR and added them to the pWPXL vector with Gibson Assembly Master Mix (NEB). We then constructed final transfer vectors by inserting target cDNA after the P2A using Gibson assembly. All constructed vectors were sequence confirmed at Gibson assembly junctions by Sanger sequencing (Genewiz) prior to use. Plasmids were grown and stored in NEB Stable *E. coli* (NEB).

### Lentiviral overexpression and flow cytometry analysis

To produce virus, we used jetPrime (Polyplus) according to the manufacturer’s instructions to transfect Lenti-X 293 T HEK cells (Takara) with lentiviral production plasmids pMD2.G and psPAX2 along with our individual transfer plasmids. pMD2.G and psPAX2 were gifts from Didier Trono (Addgene plasmid #12259; http://n2t.net/addgene:12259; RRID:Addgene_12259; Addgene plasmid #12260; http://n2t.net/addgene:12260; RRID:Addgene_12260) We collected viral media at 24 and 48 hours and concentrated using Lenti-X Concentrator (Clontech) according to the manufacturer’s instructions.

We seeded human stem cells in mTeSR medium containing γ-27632 on Matrigel-treated membrane filters as described above. We treated cells with 1x and 3x viral titer at 24 hours and 48 hours post-seeding, respectively, in order to obtain transduction efficiency of ∼10%. 1x viral treatment was performed simultaneously with the beginning of A83-01 treatment.

Two samples of each overexpression condition were performed in parallel. We harvested cells of one sample after 3 days of treatment with A83-01 and the cells of the second sample after 3 days of A83-01 followed by 42 hours of BMP4 + Activin A treatment. We analyzed each sample immediately after harvest using an LSRFortessa (BD Biosciences).

We analyzed differential OCT4:RFP to SOX2:YFP ratio distributions between CFP-positive and CFP-negative populations of each 3-day sample by calculating the Kullback-Leibler divergence in MATLAB (MathWorks). To determine differences in proportions of end fates (OCT4:RFP+/SOX2:YFP- and OCT4:RFP-/SOX2:YFP+), we manually gated ectoderm and mesendoderm populations using a custom MATLAB script and used identical gates for both CFP-positive and CFP-negative populations. We performed at least 3 biological replicates for all candidates that showed an initial phenotype except for SOX2, AHR, ARNT2, and GBX2, each of which had two replicates. Significance compared to the CFP-overexpressing negative control was determined using a two-sided *t*-test, and we controlled the FDR at 10% across the set of all candidates using the method of Benjamini and Hochberg^35^.

### Immunofluorescence

Cells were fixed with 4% formaldehyde for 15 min at room temperature. Fixed cells were treated with blocking buffer (PBS + 5% normal donkey serum + 0.3% Triton X-100) for 1 h, then overnight at 4°C with primary antibody diluted in staining buffer (PBS + 1% BSA + 0.3% Triton X-100). The following primary antibodies were used: OCT4 (1:400, Cell Signaling C30A3); SOX2 (1:400, Thermo Fisher BTJCE); SOX17 (1:100, R&D Systems AF1924); phosphorylated SMAD1/5/9 (1:200, Cell Signaling D5B10); phosphorylated-SMAD2 (1:200, Cell Signaling E8F3R); PAX6 (1:200, DSHB AB_528427); and NANOG (1:500, R&D Systems AF1997). The PAX6 antibody was deposited to the DSHB by Kawakami, A. (DSHB Hybridoma Product PAX6). After overnight incubation, samples were washed three times with PBS, then secondary antibodies diluted in staining buffer were added. We used the following secondary antibodies all at a dilution of 1:1000: donkey anti-rabbit Alexa 568 (Thermo Fisher), donkey anti-rat Alexa 488 (Thermo Fisher), and donkey anti-mouse Alexa 647 (Thermo Fisher). We incubated with a 300 nM DAPI (Thermo Fisher) solution in PBS for 5 minutes to visualize DNA. For analysis of the resulting images, we used CellProfiler 3.1.8^36^ to segment well-separated nuclei for samples where automated segmentation performed well (those shown in Figure 2), and we used manual segmentation for more challenging images (those shown in Figures 1, S1, and S6).

### ATAC-seq analysis

Reads were trimmed using NGmerge 0.2_dev in adapter removal mode with minimum overlap (- e flag) set to 20 to remove any remaining adapter sequence. Reads were aligned to the hg38 build of the human genome using bowtie2 2.2.9 using the --very-sensitive preset and with a maximum fragment size of 2000, then collated with samtools 1.9. Duplicate fragments were removed with picard 2.8.0. Peaks were called with MACS2 2.1.1 in callpeak -f BAMPE mode. Differentially accessible peaks were identified using the Bioconductor package DiffBind 2.12.0 in R 3.6.1. Peaks were annotated by genomic region type using ChIPSeeker 1.20.0.

For differential accessibility analysis with DiffBind and DESeq2, we used a design matrix with an “sample” column, which indicated the well from which the cells had been sorted (since each pair of pre- and post-competence-loss samples was derived from a single population sorted by FACS), and a “competenceloss” column, which was 1 for the post-competence-loss population and 0 for the pre-competence-loss and mesendoderm-derived populations. Thus, we identified regions that changed specifically with competence loss while controlling for original sample identity.

The primary DNA sequences of differentially accessible peaks were retrieved from Ensembl and examined for motifs using MEME-ChIP 5.0.3. ATAC-seq read depth was modeled as a function of known motif presence using chromVAR 1.4.1. Significant motif matches were identified with FIMO 5.0.3. For the gene regulatory network, possible associations between genomic regions and target genes were identified using CisMapper 5.0.5. The full list of human TFs and the motifs for each TF were extracted from the list in Lambert *et al.*^37^. Mutual information between pairs of motifs was calculated with a custom python script.

Enrichment of functions of genes near differentially accessible genomic regions was performed using the web interface of GREAT^31^ in June 2019.

### RNA-seq analysis

Reads were pseudoaligned using kallisto 0.45.1 to transcripts from the human genome build hg38. Differentially expressed genes were identified using DESeq2 1.24.0 on R 3.6.1. For analysis with DESeq2 when comparing pre- and post-competence-loss populations, we used a design matrix with an “sample” column, which indicated the well from which the cells had been sorted and a “population” column, which indicated the pre- or post-competence-loss state. For comparison of pre-competence-loss and mesendoderm-derived populations, our design matrix contained sequencing batch and pre-competence-loss or mesendoderm-derived population identity.

For clustering the motifs of differentially expressed transcription factors, similarity between each pair of motifs was quantified as the Kullback-Leibler divergence of the product of the two motifs from a reference distribution, which was the product of two uniform motifs (0.25 probability for each base at each position). Motif alignment was performed by calculating the aforementioned divergence at each possible offset and using the maximum value obtained at any offset. This calculation was performed using a custom python script. The linkage was computed using the scipy.cluster.hierarchy.linkage function from scipy 1.3.0 with the “average” clustering method and the “braycurtis” distance.

ChIP-seq target set enrichments were calculated using the Enrichr^38^ web interface in July 2019. GO term enrichment was calculated using the PANTHER^39^ web interface in July 2019.

### Overexpression candidate selection

We selected TFs by incorporating information from both RNA-seq and ATAC-seq analyses. We began with all TFs that were differentially expressed between the pre- and post-competence-loss populations (q < 0.05 with DESeq2). A gene was considered to be a TFs if it was so annotated in Lambert *et al.*^37^. We then limited this list only to those that were expressed in a lineage-specific pattern and had above-background expression levels in at least one of the three populations. We defined genes with a lineage-specific expression pattern as those genes that (1) were differentially expressed between pre- and post-competence-loss populations and (2) either were not differentially expressed from the pre-competence-loss to mesendoderm populations or were differentially expressed in the opposite direction (upregulated from pre- to post- and downregulated from pre- to mesendoderm, or vice versa). By these criteria, 23 TFs were specifically upregulated with competence loss and 32 were specifically downregulated with competence loss. We also added select paralogs of the TFs that passed our expression pattern cutoffs: POU6F1, GRHL1, POU2F3, and FOXJ2, along with the OCT4 and SOX2 DNA binding domains. We further restricted the list to those candidates that had a known, high-quality DNA binding motif that appeared in either the DiffBind/MEME-ChIP or chromVAR analyses of our data. We also added four TFs (OTX2, JUNB, ZSCAN23, and GSC) whose motifs appeared in our ATAC-seq analyses but did not pass our differential expression cutoffs. We excluded ZEB1 and ZEB2 as candidates because their size precluded delivery by lentivirus. We also eliminated 10 TFs for which a clone was not readily accessible to us, either from the Harvard PlasmID database, Addgene, or genes that had previously been cloned from cDNA in our lab. We note that one candidate that was tested before all RNA-seq analysis was complete, MBNL2, missed significance cutoffs in the final analysis but is nevertheless included for completeness. After adding three candidates based on the literature (NRF2, ZNF521, and ID2), we were left with 36 candidates in total.

### Epifluorescence imaging of fixed samples

Samples were imaged on a Zeiss AxioObserver Z1 inverted microscope using Zeiss 10x and 20x plan apo objectives (NA 1.3) using the appropriate filter sets. Images were acquired using an Orca-Flash 4.0 CMOS camera (Hamamatsu). The 43 HE DsRed/46 HE YFP/47 HE CFP/49 DAPI/50 Cy5 filter sets from Zeiss were used. The microscope was controlled using the ZEN software.

### Live cell time lapse imaging

Samples were imaged on a Zeiss AxioObserver Z1 inverted microscope using a Zeiss 20x plan apo objective (NA 0.8) using the appropriate filter sets and a Hamamatsu ImagEM EMCCD camera. Cells were maintained in a 37 degree incubation chamber at 5% CO_2_. Cells were imaged every 15 minutes. Focus was maintained using a combination of Zeiss Definite Focus and, using a custom script in MicroManager 2.0 beta^40^, software autofocus adjustments every hour to compensate for slight movement of the membrane. For maximum accuracy, cells in this time lapse were tracked manually, and the tracks were analyzed with a custom python script that performed illumination profile correction and background subtraction.

### Confocal imaging

For Figure 1 and S1, cells were imaged on a Leica inverted microscope with a Zeiss 20x objective (NA 0.8) with the appropriate filter sets. Detection was performed with photomultiplier tubes (for detection of Alexa 488 and Alexa 647) and a Leica HyD Photon Counter (for Alexa 568). For Figure S6, cells were imaged on a Zeiss LSM 880 with Airyscan using a Zeiss 20x objective (NA 0.8). Detection was performed with photomultiplier tubes (Alexa 568 and Alexa 647) and a GaAsP detector (CFP and Alexa 488).

### p(mesendoderm|OSR) curve fitting and location inference

For the initial *p*(mesendoderm|OSR) curve fitting to the single cell data extracted from the time lapse, we fit a two-parameter sigmoid function 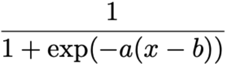 to the data using scipy.optimize.curve_fit to minimize the squared difference between data and prediction. We used the learned sigmoid shape parameter, a, for all subsequent *p*(mesendoderm|OSR) inference. To infer *p*(mesendoderm|OSR) for a given population, we fit the location parameter, b, by minimizing the squared difference between the observed final mesendoderm fate proportion and the mesendoderm proportion predicted by *p*(mesendoderm|OSR) at varying locations, b, given the observed *p*(OSR|*t*).

### Ethical compliance

We used hESCs in accordance with approvals by Harvard University IRB (protocol #IRB18-0665) and Harvard University ESCRO (protocol E00065).

## Data availability

The RNA-seq and ATAC-seq datasets generated in this study will be made available in the NCBI GEO repository and the NCBI SRA by the time of publication, or in advance of that date upon reasonable request. All other data are available upon reasonable request.

## Code availability

All custom code used in this study is available from the authors upon reasonable request.

## Author contributions

J.R.V. and S.R. conceived the project. J.R.V., R.H., S.K., and S.R. designed the experiments. J.R.V., R.H., S.K., and D.V. conducted the experiments. J.R.V. and R.H. analyzed the data. J.R.V., R.H., S.K., D.V., R.E.K., and S.R. wrote the manuscript.

## Acknowledgments

We thank the staff of the Bauer Core at Harvard University for their work on the RNA sequencing used in this manuscript as well as for their expertise and assistance with flow cytometry and FACS. We thank the Massachusetts General Hospital sequencing core for their work on the ATAC-seq data used in this manuscript, Manashree Damle for bioinformatics help, the Harvard Physics/SEAS Instructional Machine Shop for making the stainless-steel washers used in the live cell imaging in this study, and the Harvard Center for Biological Imaging for the use of their equipment. We thank Andrew Murray, Sean Eddy, and all of the members of the Ramanathan Lab for their helpful comments. J.R.V. was funded by The Fannie and John Hertz Foundation, the National Science Foundation Graduate Research Fellowship Program, and the NSF-Simons Center for Mathematical and Statistical Analysis of Biology at Harvard, award #1764269. Some confocal imaging was conducted on an instrument provided by the Harvard MRSEC (DMR-1420570). This work was supported in part by NIH R01GM131105-01.

---

**Figure S1:**
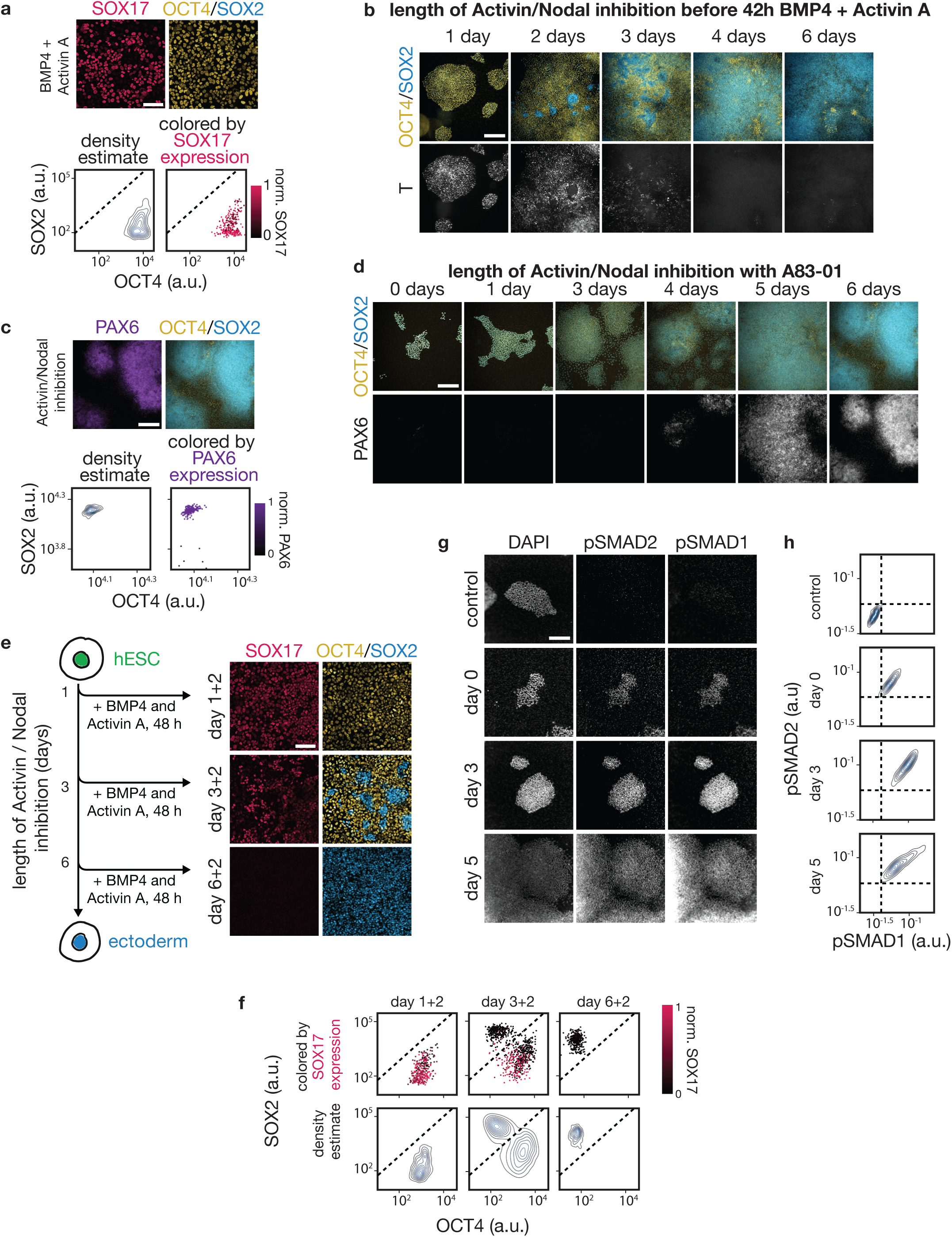
Differentiating stem cells produce mesendoderm-derived and ectoderm-derived cell types while maintaining the ability to receive BMP4 and Activin signals. **a,** Top, confocal microscopy image of immunostained hESC colony after 48 hours of BMP4 + Activin A; SOX17 (pink), OCT4 (yellow), SOX2 (blue). These cells largely adopt SOX17^+^ endoderm fates and are uniformly OCT4^+^ SOX2^-^. Scale bar = 100 μm. Bottom Left, contour plot of cell density as a function of OCT4 and SOX2 levels in individual cells; Bottom Right, same plot as left displayed as scatter plot with individual cells colored based on the level of endodermal marker SOX17. **b,** Images of hESC colonies immunostained for OCT4, SOX2, and mesendoderm marker T after 1-6 days of Activin/NODAL inhibition followed by 42h of BMP4 + Activin A stimulation. T^-^ OCT4^-^ SOX2^+^ ectoderm cells first appear at 2 days and are widespread by 4 days of Activin/NODAL inhibition. The spatial structure seen here largely appears after BMP4 and Activin A signal induction and is likely due to a combination of local density impacts on ectoderm-directed differentiation rate and homophilic interactions between cells. Scale bar = 300 μm. **c,** Top, epifluorescence microscopy image of an hESC colony immunostained for the neurectoderm marker PAX6 (purple), OCT4 (yellow), and SOX2 (blue) after 6 days of Activin/NODAL inhibition. These cells largely adopt PAX6^+^ OCT4^-^ SOX2^+^ neurectoderm fates. Below, contour plot of cell density as a function of OCT4 and SOX2 levels in individual cells (left), same plot as left displayed as scatter plot with individual cells colored based on the level of PAX6 (right). Scale bar = 300 μm. **d,** Images of hESC colonies immunostained for OCT4, SOX2, and the neurectoderm marker PAX6 after 0-6 days of Activin/NODAL inhibition. PAX6 expression is first seen after 4 days and is widespread by 5 days. Scale bar = 300 μm. **e,** hESCs were exposed to a pretreatment of Activin/NODAL inhibition for 1, 3, or 6 days before BMP4 + Activin A signal exposure (left). Confocal microscopy images of immunostained hESCs at the end of the experiment after 48h of BMP4 + Activin A treatment (right). Increasing duration of Activin/NODAL inhibition reduced the population’s competence to produce SOX17^+^ endoderm and, more broadly, OCT4^+^ (yellow) SOX2^-^ (blue) mesendoderm-derived cell types. Scale bar = 100 μm. **f,** Scatter plot (top) and density contour plot (bottom) of cells from E, as a function of their OCT4 and SOX2 levels. Cells in scatter plot colored by level of SOX17. **g,** Immunofluorescence images acquired via epifluorescence microscopy showing DAPI, phosphorylated SMAD1, and phosphorylated SMAD2 levels in hESC colonies upon stimulation with BMP4 + Activin for 45 mins after 0, 3, or 5 days of Activin/NODAL inhibition. Unstimulated control shown in top row. All stimulated cells at all time points displayed nuclear-localized pSMAD1 and pSMAD2 upon BMP4 + Activin A exposure despite the loss of mesendoderm competence that occurs during this window. Scale bar = 300 μm. **h,** Density contour plots of cells treated as in G, as a function of pSMAD2/3 and pSMAD1/5 levels determined by antibody staining. Dotted lines drawn for reference.

**Figure S2:**
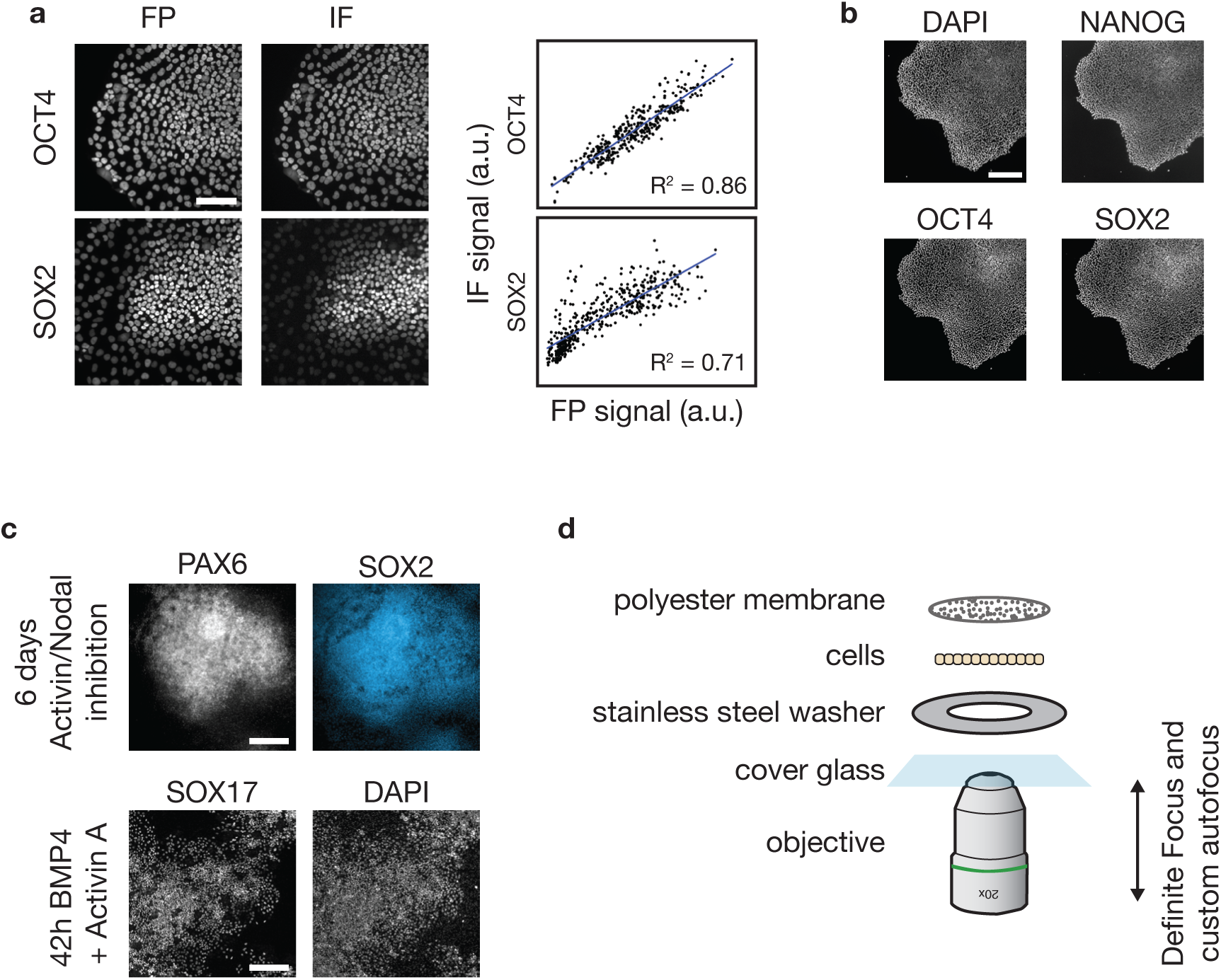
Monitoring OCT4:RFP and SOX2:YFP accurately reflects OCT4 and SOX2 levels without perturbing cell potential or behavior. **a,** OCT4 and SOX2 levels as determined by immunofluorescence (IF) and by OCT4:RFP and SOX2:YFP fluorescence (FP). Scale bar = 100 μm. Blue line, best-fit linear regression. **b,** Immunofluoresence images of H1 OCT4:RFP SOX2:YFP double reporter cells stained for NANOG, OCT4, and SOX2. Nuclei (DAPI) are shown for reference. These cells can maintain pluripotency indefinitely. Scale bar = 300 μm. **c,** Immunofluorescence images of H1 OCT4:RFP SOX2:YFP double reporter cells that were (top) stained for PAX6 and SOX2 after 6 days of Activin/NODAL inhibition by A83-01 or (bottom) stained for SOX17 after 42h of BMP4 and Activin A with nuclei (DAPI) shown for reference. This line’s ability to form the germ layers is unaffected. Scale bar = 300 μm. **d,** Schematic of the live-cell imaging apparatus constructed for this study. Cells were grown on the bottom of a permeable membrane glued to a thin stainless-steel washer and imaged from below with an epifluorescence microscope. Focus was maintained with a combination of Zeiss Definite Focus and periodic software autofocus adjustments.

**Figure S3:**
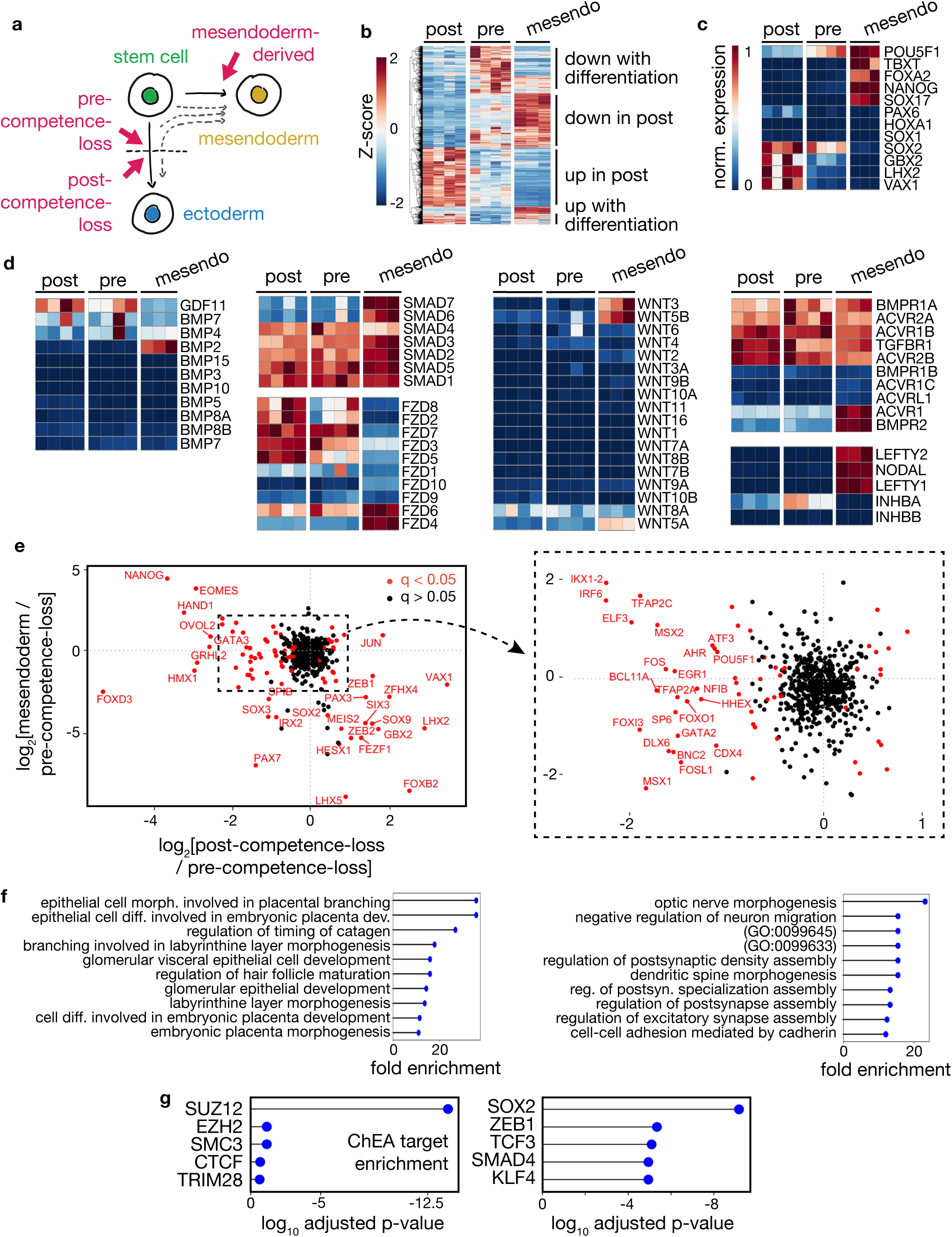
Analysis of gene expression changes that coincide with mesendoderm competence loss. **a,** Based on the OCT4:RFP to SOX2:YFP ratio, pre- and post-competence-loss cells were isolated using FACS for RNA-seq and ATAC-seq. Mesendoderm-derived cells (hESCs stimulated with 40 h of BMP4 + Activin A) were also assayed as an outgroup to allow for the identification of ectoderm lineage-specific changes. **b,** Heatmap showing normalized gene expression changes for genes that display significant differential expression between at least two of the three assayed populations in (a). **c,** Heatmap showing normalized expression of selected marker genes whose expression is associated with the early germ layer lineages. Gene expression was normalized by the maximal expression observed in each row or 10 TPM, whichever was greater. **d,** Heatmap showing normalized expression of selected signaling molecules, transducers, and receptors. Gene expression was normalized by the maximal expression observed in each row or 10 TPM, whichever was greater. No significant expression changes between the pre- and post-competence loss populations were observed in these groups. **e,** Scatterplot showing log2 fold change for all transcription factors between the (x-axis) post- and pre-competence-loss populations and the (y-axis) mesendoderm and pre-competence-loss populations. TFs that are significantly differentially expressed between the pre- and post-competence-loss populations are shown in red. TFs in the second and fourth quadrants of this plot have lineage-specific expression patterns. **f,** Ten most significantly enriched GO terms for gene expression sets from this study as calculated by PANTHER. Left, enriched terms for genes that are downregulated upon competence loss in a lineage-specific pattern. Right, same for genes that are upregulated. **g,** Left, most significant ChIP-seq target set enrichments as reported by Enrichr for genes whose expression was upregulated from pre-competence-loss to post-competence-loss cells but not significantly upregulated from the pre-competence-loss to mesendoderm-derived populations. Right, same for genes that were downregulated from pre- to post-competence-loss but not downregulated from pre-competence loss to mesendoderm populations.

**Figure S4:**
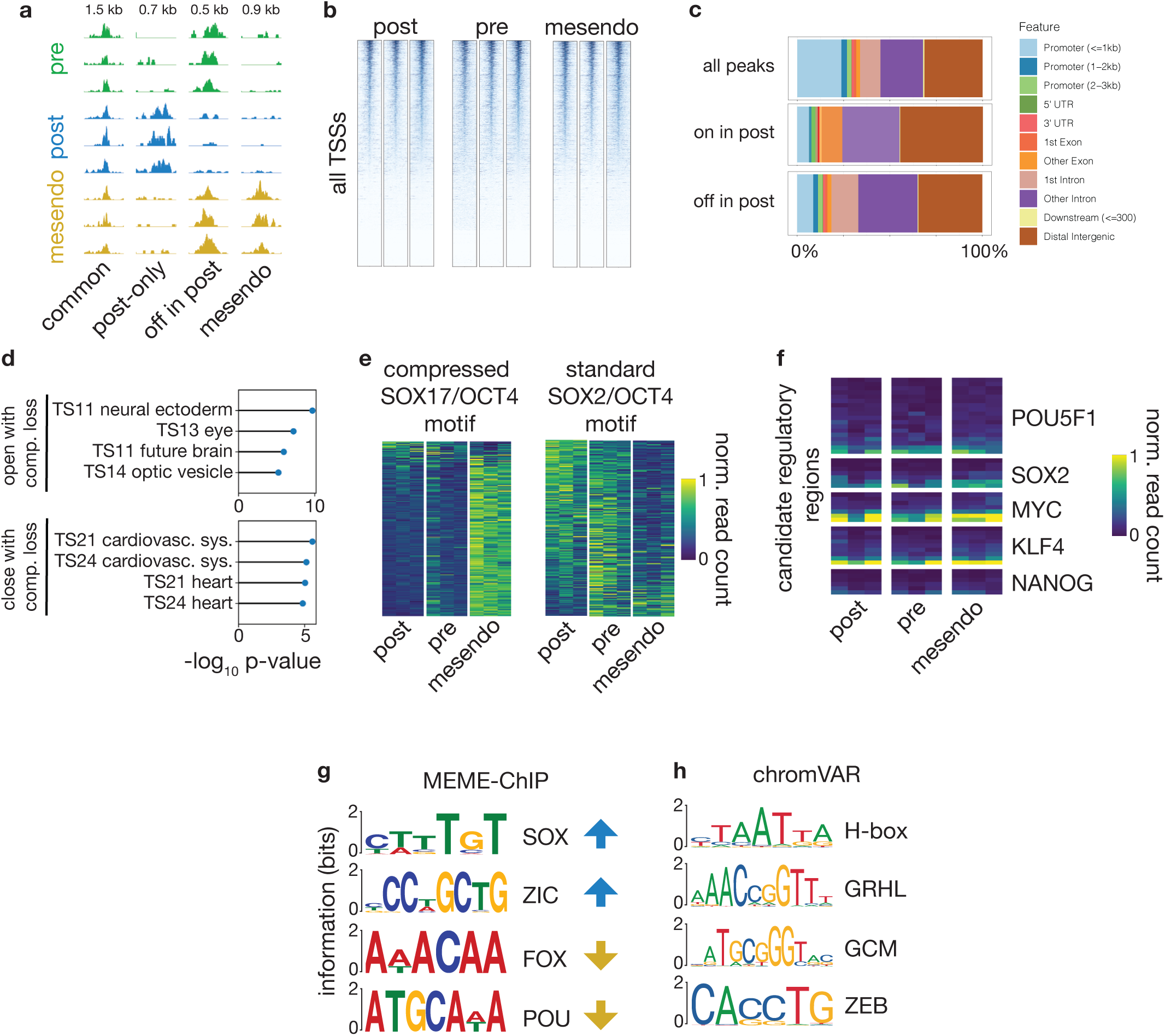
Analysis of chromatin accessibility changes that coincide with mesendoderm competence loss. **a,** Normalized ATAC-seq read depth at example loci with the indicated accessibility patterns across pre-competence-loss, post-competence-loss, and mesendoderm-derived populations. Text at top indicates the length of the region shown. **b,** ATAC-seq read depth at all transcription start sites. Each heatmap block represents all transcription start sites (TSSs) in a single biological replicate for the indicated condition. Each row shows a heatmap representation of read depth in 2kb windows centered on each TSS. Rows are sorted for each replicate individually by mean read depth across the window. **c,** Genomic feature type of the indicated ATAC-seq peaks as annotated by ChIPSeeker. Annotations are shown for all detected peaks across all samples, for peaks that are significantly more accessible in post-competence-loss cells, and for peaks that are significantly less accessible in post-competence-loss cells. Most differentially accessible sites are located in distal intergenic or intronic regions. **d,** Overlap enrichment of mouse homologs of the differentially expressed gene sets from this study with developmental stage gene expression sets from the Mouse Genome Informatics database as calculated by the software GREAT. Top, the four most enriched gene expression set matches for genes that are upregulated in a lineage-specific manner upon competence loss. Bottom, the same for genes that are downregulated. **e,** Heatmap showing row-normalized mean ATAC-seq depth in 500 bp regions centered on each peak that shows significant changes in accessibility and which contains the indicated motif. Left, the regions that contain a compressed SOX17/OCT4 motif associated with endoderm occupancy. Right, the same for regions that contain a standard SOX2/OCT4 motif associated with pluripotency occupancy. **f,** Heatmap showing normalized ATAC-seq read depth in all ENCODE-annotated candidate regulatory regions for the indicated pluripotency-associated genes. None of these regions show significant differences between pre- and post-competence-loss cells. **g,** Selected motifs with significant enrichment in genomic regions that had a significant increase (blue arrows) or decrease (yellow arrows) from the pre-competence-loss to the post-competence-loss populations as discovered using MEME-ChIP. Motifs are labeled by the TF family/group to which they correspond. E-values: SOX, 3.7e-128; ZIC, 8.3e-033; FOX, 1.8e-02; POU, 2.9e-003. **h,** Top 4 nonredundant motifs that best explain the observed changes in ATAC-seq read depth between pre-competence-loss and post-competence-loss populations, as calculated using chromVAR. p-values: homeobox (VAX2), 9.8E-05; GCM (GCM1), 3.2E-4; GRHL (GRHL1), 6.6E-4; ZEB (ZEB2), 6.9E-4.

**Figure S5:**
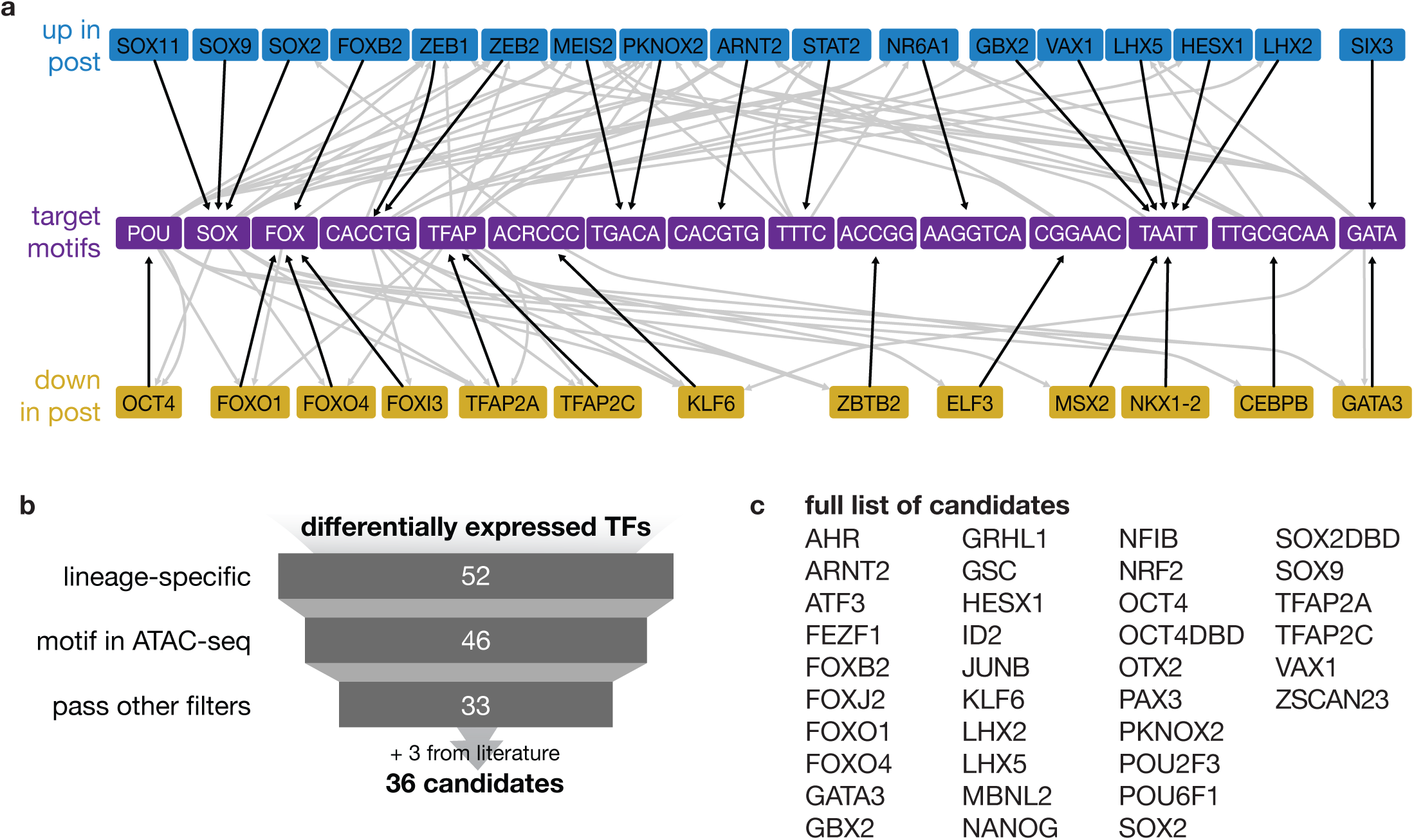
Gene expression and chromatin accessibility signatures reveal a core set of TFs that, when perturbed, could modulate competence. **a,** Candidate gene regulatory network that governs mesendoderm competence. TFs that are upregulated (top row, blue) or downregulated (bottom row, yellow) specifically between pre- and post-competence-loss populations are connected to their DNA binding motifs by black arrows. Each motif is connected to possible regulatory targets by gray arrows according to the following criteria (1) there is a significant match to that motif in an ATAC-seq peak that is (2) predicted to be associated with the target gene via the CisMapper software. **b,** Schematic depicting the candidate selection process (see methods for full details). **c,** Full list of the 36 TFs identified as candidates for perturbation to affect mesendoderm competence

**Figure S6:**
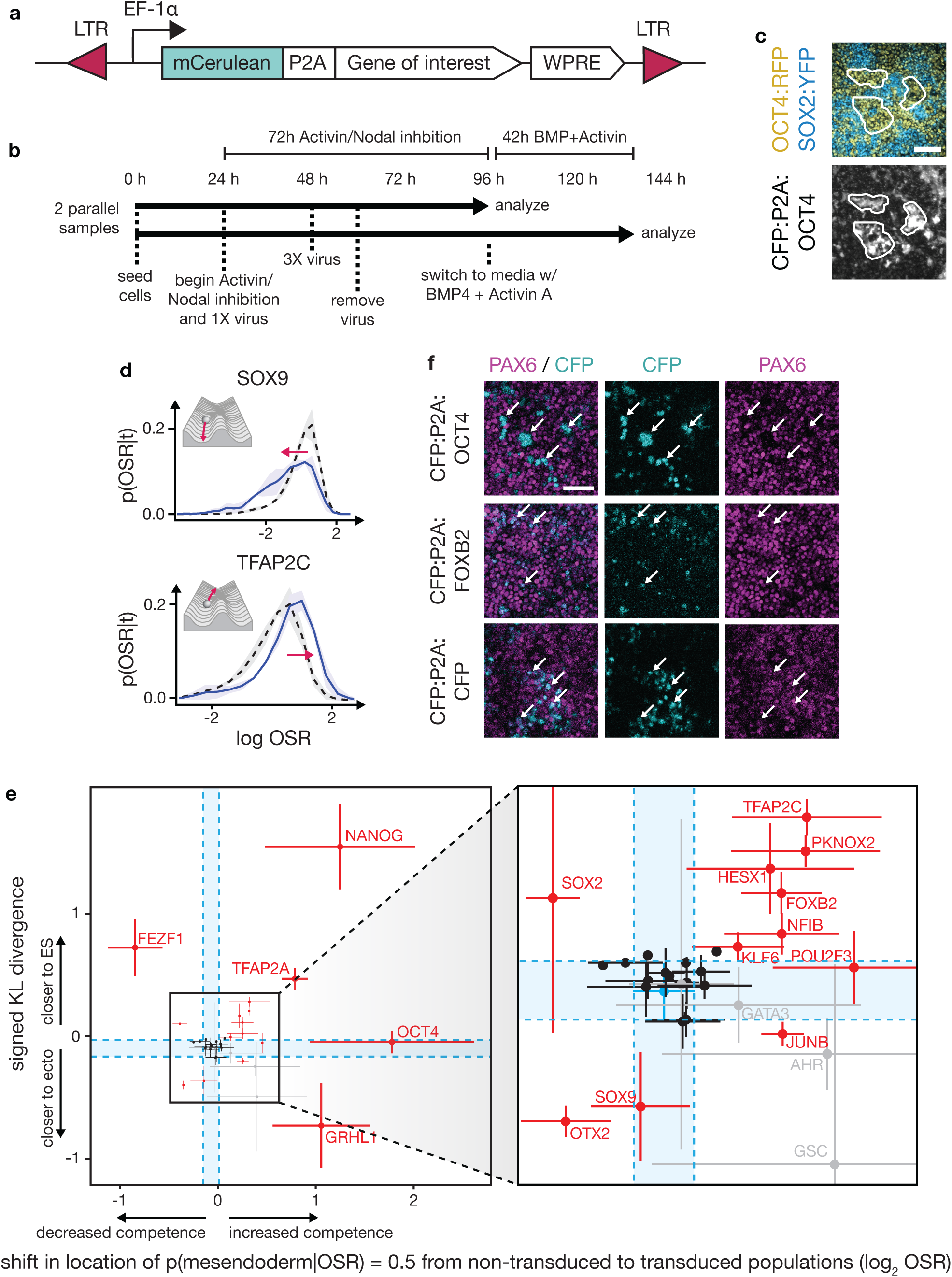
Analysis of lineage bias and differentiation dynamics in cells perturbed by candidate TFs. **a,** Schematic of the cassette transduced via lentiviral vector. Under the control of the EF1alpha promoter, we placed the cyan fluorescent protein mCerulean fused in frame to a P2A ribosomal skip sequence and the gene of interest. A Woodchuck Hepatitis Virus Posttranscriptional Regulatory Element (WPRE) was included to improve transcript stability. The entire sequence was flanked by the Long Terminal Repeat regions (LTR) that the viral transposases recognize to effect integration into the genome. **b,** Schematic of experimental protocol. 24h after seeding in pluripotency maintenance conditions, parallel samples of hESCs were subjected to simultaneous Activin/Nodal inhibition and, for the first 36h, lentiviral transduction of the gene of interest. After 72h of Activin/Nodal inhibition, one membrane was analyzed by flow cytometry while the other was stimulated with 42h of BMP4 + Activin A before analysis by flow cytometry. **c,** Epifluorescent microscopy images of a cell colony after Activin/Nodal inhibition and subsequent BMP4 + Activin A stimulation. Cells that were transduced with a CFP:P2A:OCT4 expression cassette expressed CFP and preferentially adopted a OCT4:RFP^+^ SOX2:YFP^-^ mesendoderm fate. White regions indicate selected clusters of cells expressing CFP, indicating transduction. Scale bar = 100 μm. **d,** Examples of two candidates, SOX9 and TFAP2C, whose overexpression altered the distribution of OCT4:RFP/SOX2:YFP ratios of cells before the signal, p(OSR | t). OCT4:RFP and SOX2:YFP levels were normalized to an hESC sample measured in the same batch. Black dotted line, wildtype. Blue, transduced. Shaded area represents one standard deviation across biological replicates. Inset, a schematic depicting the corresponding effects on the Waddington developmental landscape. **e,** Scatterplot of the shift in p(mesendoderm|OSR) location versus a metric of disruption of p(OSR|t) for each TF tested. Disruption of initial differentiation dynamics was measured as the Kullback-Leibler divergence of the distribution of the log ratio of OCT4:RFP to SOX2:YFP of CFP^+^ cells using the same distribution from CFP^-^ cells as a reference; this KL-divergence was then reported with a positive sign if the mean OSR of the transduced cells was higher than in the non-transduced cells and a negative sign if not. Candidates that were significantly different from the CFP negative control (FDR < 0.1) along either dimension are shown in red. Candidates whose effects were not consistent across replicates are shown in gray. CFP:P2A:CFP is shown in blue. Blue dotted lines indicate CFP:P2A:CFP overexpression outcomes, shown as the mean ± one standard deviation of n=4 biological replicates, for each axis. OCT1, POU6F1, PAX3 were tested but are excluded from this plot because their overexpression caused premature downregulation of both OCT4:RFP and SOX2:YFP, indicating possible differentiation to an alternative fate. Interestingly, exogenous OCT4 overexpression does not impact the dynamics of endogenous OCT4:RFP and SOX2:YFP, as evidenced by the location of the OCT4 point near zero on the y-axis. **f,** Immunofluorescence images showing PAX6 (magenta) and CFP (cyan) after 6 days of neurectoderm-directed differentiation using inhibitors of both Activin/NODAL and BMP. Top row, exogenous expression of CFP:P2A:OCT4; Middle row, CFP:P2A:FOXB2; Bottom row, CFP:P2A:CFP. White arrows indicate select CFP-expressing cells. Look-up tables for the channels in each condition are scaled individually because variable membrane positioning in the well created spurious changes in overall apparent signal intensity. Scale bar = 50 μm. OCT4 overexpression prevents PAX6 induction, while FOXB2 or CFP overexpression does not disrupt normal PAX6 induction.

